# Characterizing the expression patterns of short-day flowering genes, the manipulation of their expression using red-far red wavelengths, and the resulting phenotype/ chemotype in *Cannabis sativa*

**DOI:** 10.1101/2024.03.28.587211

**Authors:** Samuel Haiden

## Abstract

Cannabis is a short day (SD) plant whose inflorescences produce valuable medicines. By studying the mechanisms of photoperiodic flowering initiation in cannabis, we may develop new methods of improving cultivation. In SD plants, CONSTANS, which is enhanced by CDL gated proteins that are expressed during short days, acts as a bifunctional regulator of flower inhibition or genesis via FT (florigen). These genes are downstream of phytochrome, which may be activated by horticultural lighting conditions like red (R, 660nm) to far red (FR, 700+nm) wavelength ratios. This study demonstrates that gene expression can be manipulated using high and low R:FR ratios, and characterizes the expression levels of CONSTANS, FT, PRR37, and SOC1 in cannabis over the first 14 short days. It also demonstrates the resulting effect on cannabinoid production and biomass yields.

## Introduction

Cannabis sativa is generally a short day (SD) photoperiod-sensitive flowering crop, and its inflorescences are becoming a widely traded commodity due to their chemical contents, which include but are not limited to cannabinoids like THCa, CBDa, CBCa, and CBGa, as well as their derivatives and analogues. As such, understanding the mechanisms which determine the transition into the flowering stage is a critical area of cannabis biology.

The signaling cascade which initiates the photoperiodic flowering response is well characterized in long day (LD) plants^1^. In Arabidopsis, a model LD plant, CONSTANS acts as a central hub for integrating environmental signals to determine flowering time^2^. The FLOWERING LOCUS T gene is activated in leaves via the sequential activation of GIGANTEA and CONSTANS^3^, which are regulated by both phytochromes and cryptochromes^4^. Upon entering long days, the FT protein is positively regulated and translocated to the meristem, where it works with FD and SOC1 to activate floral meristem identity, which initiates the development of flowers^1,5^.

Cannabis is a short-day photoperiodic flowering plant and produces its flowers (inflorescences) when the critical day length is under 12-14 hours^6^. As such, it shares more similarity with rice in terms of the mechanisms of its photoperiodic-flowering induction pathway. Like arabidopsis, rice possesses a CONSTANS-like gene, but this gene is known as Hd1 or Heading date 1^7,8^. Unlike in Arabidopsis, Hd1 in rice is not a central hub for signal integration, but rather a bifunctional regulator which performs different functions based on the presence of different enhancer proteins^8^. These enhancer proteins-GHD7, PRR37 and DTH8-are CDL gated and differentially expressed based on photoperiod^9–11^. A homolog of PRR37 has been identified in cannabis, and it has been shown that a mutation of that gene is correlated with the autoflowering trait, or lack of response to photoperiod^12^. GHD7 has also been implicated as a predictor of yield in rice^10^.

Although Hd1 is a bifunctional regulator of photoperiodic flowering in rice, its expression changes very little during the transition to flowering: rather reversible, CDL-gated enhancer proteins activate it. As such, understanding these enhancer proteins is very important for our understanding of CDL photoperiodic-flowering regulation in plants, and the differences between SD and LD plants ^8,13^. Furthermore, no bi-functional reversible regulators like Hd1 have been identified in Arabidopsis^14^, indicating that this function is unique to SD plants including cannabis. Cannabis CONSTANS-like genes have been identified, and in fact there are at least 13 expressed CONSTANS-like genes^15^. In cannabis, florigen-like, PRR37-like, and SOC1-like genes have also been identified^12,16^. In this paper, we present experimental data which characterizes the expression patterns of these genes, their responsiveness to phytochrome via manipulation of the ratio of red to far red light, and the resulting impact on phenotype and chemotype.

The manipulation of phytochrome stationary state or PSS via adjusting the red to far-red ratios in horticultural lighting is a common practice in crop production^17^. This is because phytochrome is a signaling molecule which, when exposed to red or far-red light, undergoes a conformational change due to the excitation of the chromophore^18^. When it is irradiated by red light, which is present in high concentrations in full sunlight, it translocates to the nucleus and controls gene expression, which helps to entrain the circadian clock, informing the plant on a cellular level that it is daytime^1,7,18,19^. Shady light or dusk is rich in FR light, and so is moonlight. Either in the presence of FR light or in the darkness, phytochrome converts to its inactive state, and translocates again to the cytosol. The ratio of active (Pfr) to inactive (Pr) phytochrome is referred to as PSS^17^. Thusly, phytochrome controls not only photoperiodic response, but also the shade avoidance response, in which the plant will grow out of the shade and towards the full sunlight^20^. The improvement of cannabis cultivation by modifying spectral composition has been studied^21,22^.

In addition to characterizing the expression patterns of cannabis genes downstream of phytochrome during the transition to flowering, we also study the effect of modifying the PSS of the cannabis plant on gene expression. Furthermore, we test the hypothesis that by increasing the red to far-red ratio we may be able to increase cannabinoid production or flower yield.

## Results

**Figure 1:**
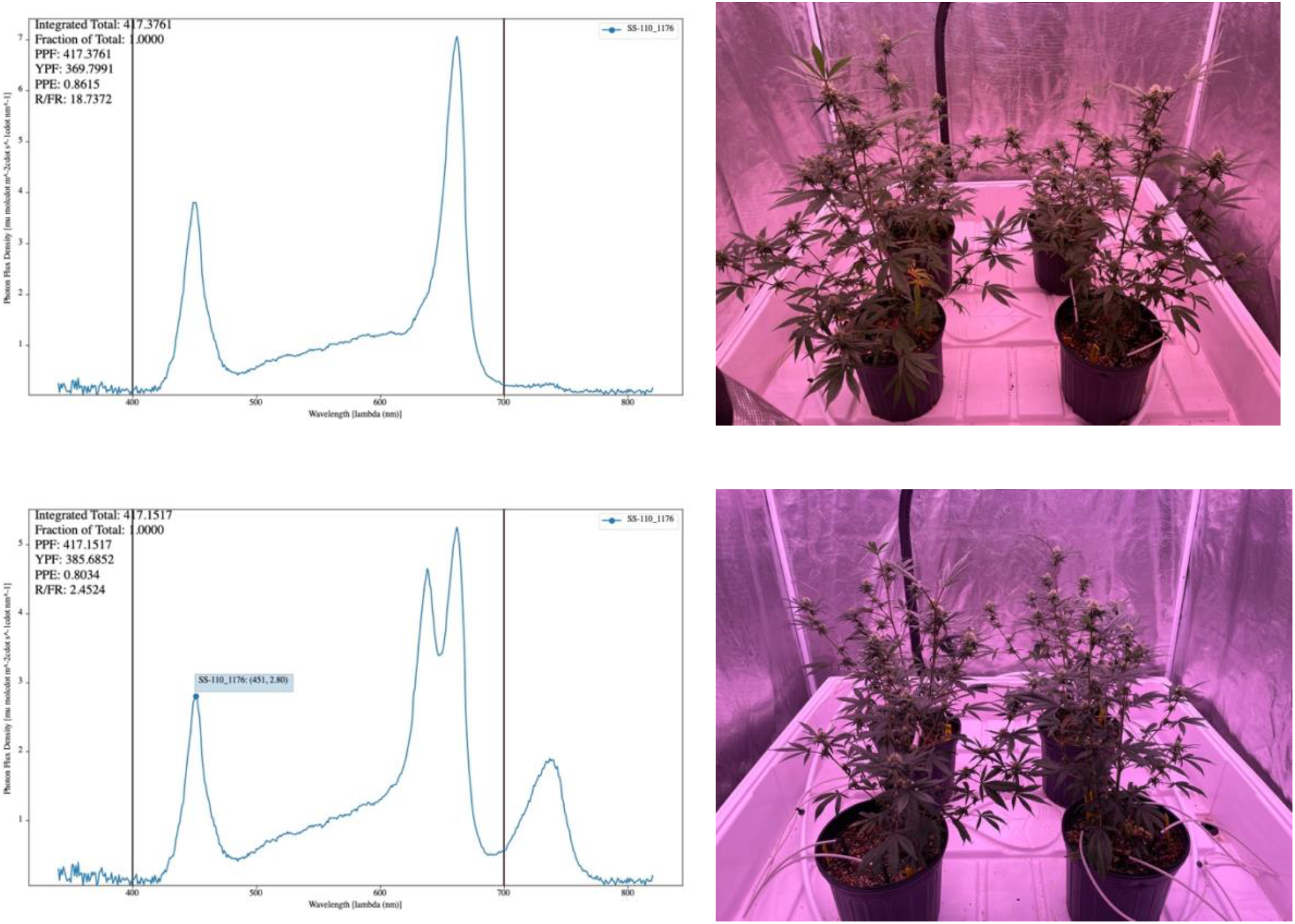
Spectroradiographs confirming the R:FR ratios generated by the light fixtures. (A) High R:FR conditions show a large peak at 660nm and no peak in the 700nm+ (FR) range. The light appears more pink. (B) Low R:FR conditions show a large peak at 660nm as well as a large peak in the FR 700nm+ range. The light appears more purple. In both conditions, PPFD and B:R ratios are normalized. Table 1 displays average treatment values and standard deviations from 5 measurement zones.

**Table 1.**

|  | High | StErr |  | Low | StErr |
| --- | --- | --- | --- | --- | --- |
| Avg RFR | 22.10 | 2.71 | Avg RFR | 2.65 | 0.09 |
| Avg R/B | 1.91 | 0.04 | Avg R/B | 1.88 | 0.03 |
| Avg PFD | 399.34 | 6.08 | Avg PFD | 398.41 | 7.69 |

**Figure 2:**
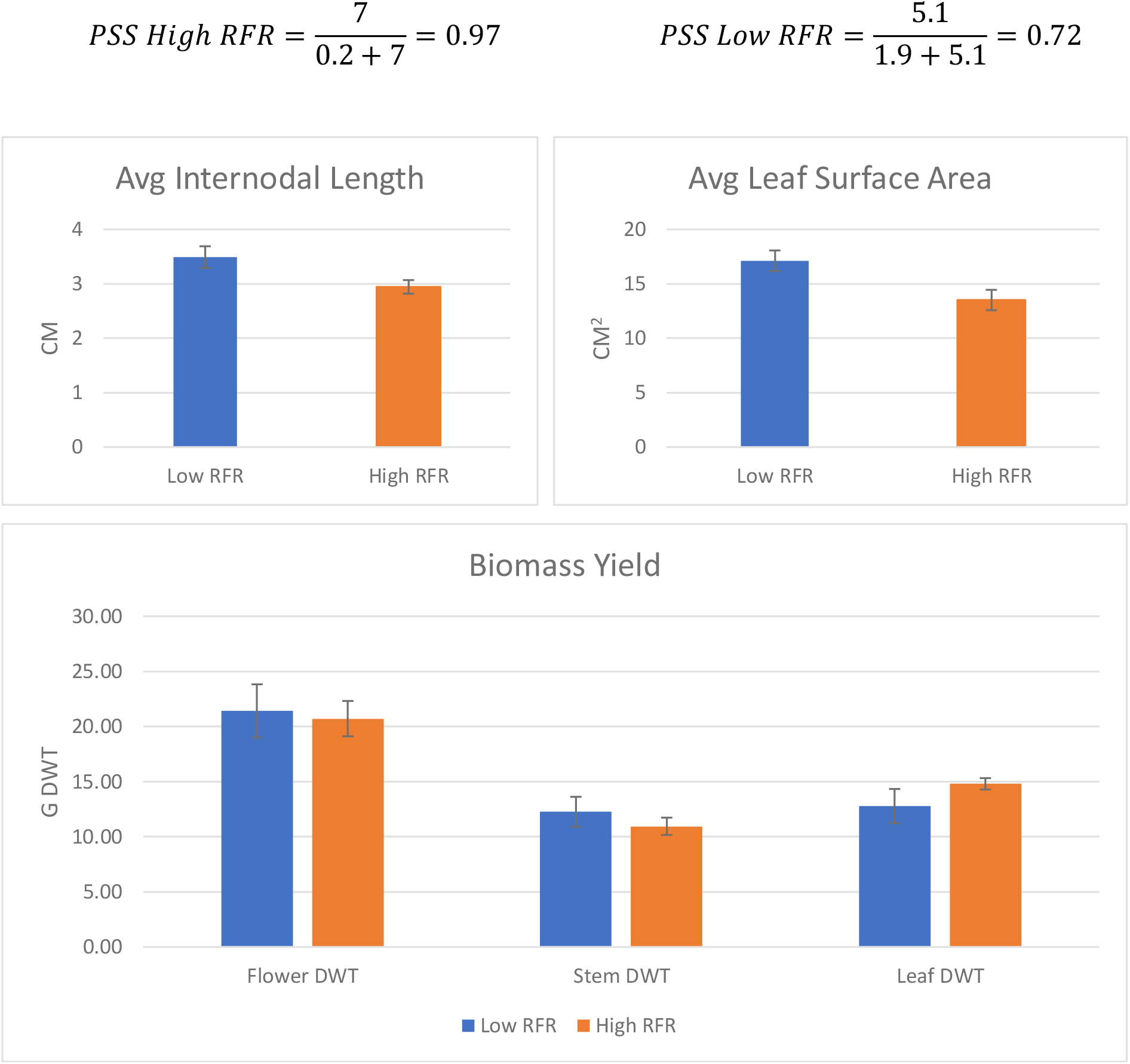
Measurements were taken on day 42 immediately prior to harvest. No change in average internodal length was observed. Leaf surface area was also unaffected. Whole plants were hung to dry for 14 days at 40% humidity and 60F until they had reached a moisture content of 10%. Tissue types were separated and weighed. No significant differences observed.

**Figure 3:**
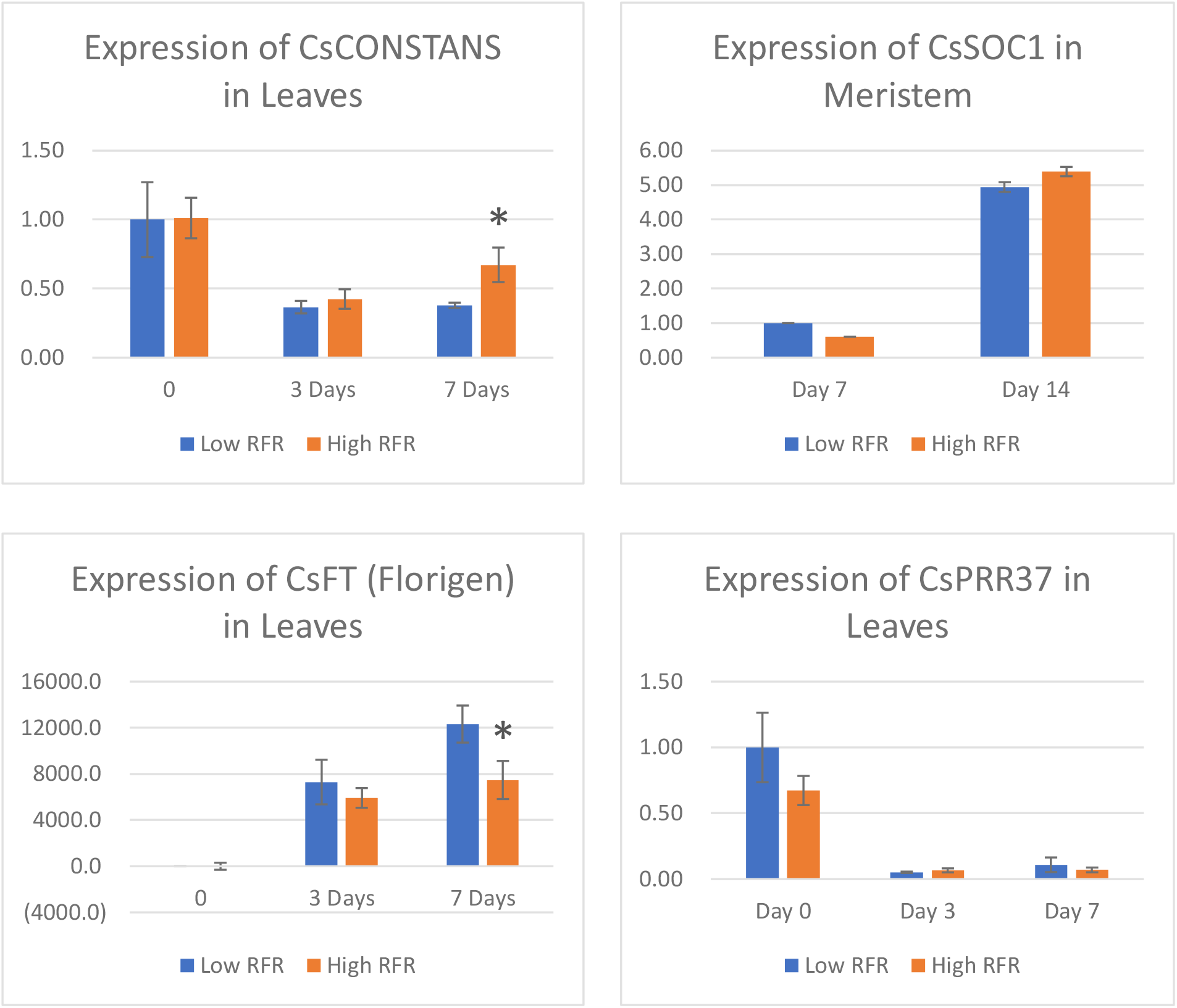
qPCR data for genes involved in the SD photoperiodic flowering initiation pathway. Day 0 refers to pre-short days, and day 3 and 7 refer to the 3^rd^ and 7^th^ short day. One star indicates a P-value below 0.01. All values calculated using the delta-delta Ct method with CsUbiquitin as a housekeeping gene. The evidence suggests that both CsFT and CsPRR37 are CDL-gated genes, where CsFT is significantly upregulated as much as 12,000 fold by the seventh short day. This is consistent with existing models for the regulation of short day flowering. CONSTANS expression generally remains stable regardless of photoperiod in SD plants, but its activity is affected by enhancer proteins like CsPRR37. PRR37 expression decreased by 20 fold after the initiation of short days, indicating it is probably a repressor of flowering, interacting with CONSTANS. As expected, SOC1 expression increases in the transitioning floral meristem during the first two weeks of short days. For each treatment group, 4 biological replicates were sampled and divided into two technical replications per plate.

**Figure 4:**
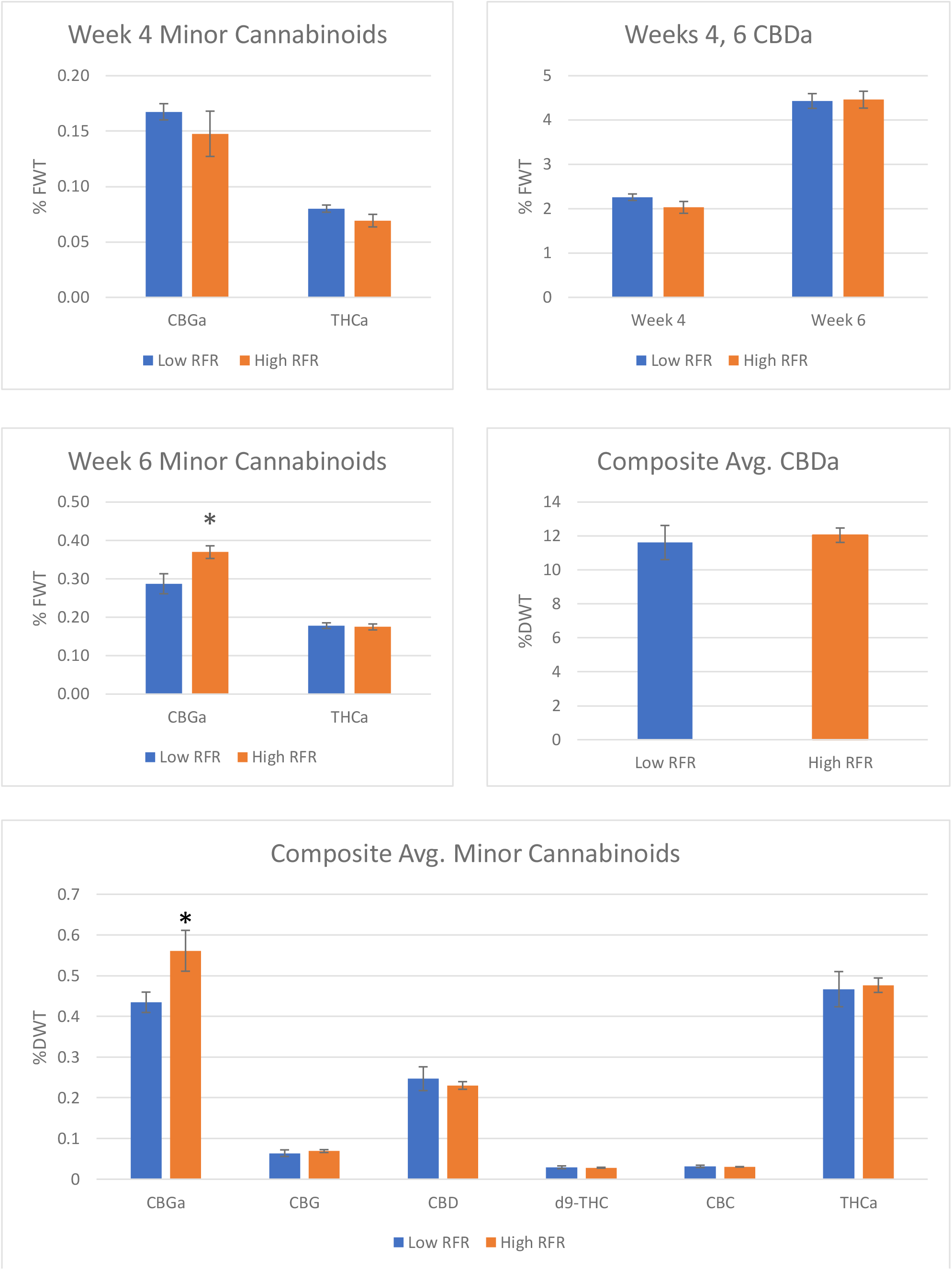
HPLC-UV results for cannabinoid content in fresh and dry flower samples. The first undamaged phytomer from the terminal shoot was harvested at days 28 and 42, snap frozen, extracted into methanol and analyzed using using the Shimadzu High-Sensitivity protocol. After two weeks of drying, each plant was milled into a composite sample, and tissue was collected from each plant composite. For each round of sampling, 4 biological replicates were sampled. 100mg of tissue was collected for each sample and extracted into 10ml of methanol, filtered, diluted, and injected. Although there is no change in the end-product cannabinoids, CBGa accumulation was increased in the high R:FR group at week 6 and in the dry composite sample.

## Discussion

This study has given us the opportunity to characterize the expression patterns of four genes which are likely to be involved in the photoperiodic flowering signaling pathway. It has also given us the opportunity to see if their increased expression yields expected results, or any impact on biomass or cannabinoid yield.

The overall expression level of our CONSTANS-like candidate did decrease slightly after the initiation of short days, but the overall change was less than appx. 2.5 fold. Interestingly, the high R:FR treatment seems to have resulted in the increased expression of CONSTANS by the seventh short day. However, given that it is likely that CONSTANS is activated by an enhancer protein, its expression level is of less consequence. This still provides evidence that increasing the red to far red ratio may have a considerable effect on gene expression, in spite of the notable fact that this change did not have any outcome on yield.

Due to the increased expression of CONSTANS, one may expect to see an increased expression level of florigen FT, but our evidence is the opposite, where FT expression was significantly decreased in the high R:FR group. Perhaps more notably, however, is that while expression of FT is approximately 0 before the initiation of short days (day 0), expression levels increase many thousand fold by the third and seventh short day. This confirms that florigen in cannabis is a CDL-gated gene, similar to HD3A (florigen) in rice. Neither SOC1 nor PRR37 expression levels were affected by the high R:FR treatment, however it is interesting to note that SOC1 levels in the transitioning flower meristem increased 5 fold between the 7^th^ and 14^th^ short day. It is also interesting to note that the expression of PRR37 decreased 20-fold after the initiation of short days, which suggests that PRR37 is not an enhancer of flowering, but is more likely to be a repressor.

Although a slightly different visual appearance was observed between the treatment groups, the data was inconsistent with visual observation. Measurement of internodal length, leaf surface area, and leaf, stem, and flower biomass showed no significant differences. However by week 6, CBGa levels in the fresh flower were more than 20% greater in the high R:FR group than the low, while end-product cannabinoid levels remained unchanged. This pattern continued through harvest and drying, where in the composite dry-weight analysis, we see an approximately 25% increase of CBGa in the high RFR treatment group, but no change in any of the other cannabinoids. This indicates that the high RFR treatment increased the expression of CBGa, but that CBGa was not the rate-limiting reactant for end-product cannabinoid synthesis in this example.

## Conclusion

In conclusion, we do not find that there is any evidence to suggest that increasing R:FR ratios in greenhouse or indoor cultivation environments will increase cannabis flower biomass or cannabinoid yields, however this study elucidated the expression patterns of multiple candidates for the genetic control of photoperiodic flowering initiation in cannabis. It appears that rice is the best model plant for understanding flowering mechanisms in cannabis, as cannabis is a short day plant. The expression patterns of CONSTANS, FT, SOC1 and PRR37 in cannabis are similar to those of their rice homologs, where FT and PRR37 are CDL-gated, expression of CONSTANS remains stable, and expression of SOC1 increases in the developing floral meristem. Although their increase or decrease in expression levels of these genes had no effect on biomass or cannabinoid yield, it is interesting to note that the expression of CONSTANS and FT were affected by high R:FR treatment. Furthermore, it is interesting to note that CBGa accumulation was increased in the high R:FR treatment group, in spite of the fact that end-product cannabinoid accumulation was unaffected. More study is needed on the mechanisms of the initiation of flowering in cannabis, as we have not yet identified key enhancers of CONSTANS or positive regulators of flowering.

## Methods

1. Plant Growth:
  a. Plants were grown inside of Gorrilla Grow grow tent structures, each outfitted with batch-tank fertigation. Humidity and temperature were controlled via high-output ventilation fans and thermostats. Cuttings were stuck in Rockwool using Clonex, allowed to root for two weeks under HPS lighting, and then planted in #900 blow molded nursery pots in ProMix BX-25 supplemented with 12g/ gallon Osmocote 15-9-12. Cuttings were then placed into grow tents and allowed to grow vegetatively for another two weeks before switching to flower. Plants were fed via drip fertigation CLF alternating between Jack’s 15-30-15 and the other one.
2. Design of Spectral Conditions:
  a. Adaptiiv ATG-1000 Liquid-Cooled LED light fixtures were used to create the spectral conditions for each treatment. Blue, Red, Far Red, and White illuminance was adjusted to achieve the target R:FR ratio while normalizing B:R ratios and PPFD. The resulting spectral composition was measured using an Apogee SS-110 Spectroradiometer, and a composite of 5 measurements was taken per treatment area to determine the variability.
3. Sampling:
  a. Leaf samples were taken immediately before switching to short days (day 0), and the follwing day was designated “Short Day 1”. The fifth, fourth, third, second and first fully expanded leaf from the terminal meristem was selected for each consecutive sampling. Leaf samples taken again at SD 1, 3, 7, and 14, at EOD appx 2 hours before lights out. Transitioning floral meristem samples taken at days 7 and 14 from the 2^nd^ and 1^st^ most dominant meristem respectively. Flower samples for RNA taken at SD 28. Flower samples taken for FWT cannabinoids from remaining undamaged terminal phytomers at SD 28 and 42. Final cannabinoid sample taken from dry composite of each plant.
4. HPLC-UV:
  a. For each analysis, 100mg of ground flower tissue was extracted into 10ml methanol, filtered, diluted and analyzed using Shimadzu Hemp Analyzer High Sensitivity Package according to manufacturer’s protocol.
5. Internodal Length, Leaf SA and Biomass
  a. The 3^rd^ branch from the terminal meristem was removed and the 1^st^, 2^nd^ and 3^rd^ leaves/ internodes were measured. Leaf SA measured using LeafScan app. Flowers dried using commercial methods and water weight calculated.
6. qPCR
  a. Ubiquitin used as housekeeping gene for all reactions. Primers designed using NCBI’s primer design tool. RNA isolated using NucleoSpin Plant and Fungi RNA Isolation Kit. cDNA synthesized using iScript Reverse transcriptase. Bio-Rad CFX-Connect Real time qPCR system was used with iTaq SYBR Green Universal Master Mix. Relative expression values calculated using delta-delta Ct method. Student’s t-tests were used to establish statistical significance.

## Notes

### Competing Interest Statement

The authors have declared no competing interest.

## Sources Cited

1. Teotia, S. & Tang, G. To Bloom or Not to Bloom: Role of MicroRNAs in Plant Flowering. Molecular Plant 8, 359–377 (2015).

2. Shim, J. S., Kubota, A. & Imaizumi, T. Circadian Clock and Photoperiodic Flowering in Arabidopsis: CONSTANS Is a Hub for Signal Integration. Plant Physiol. 173, 5–15 (2017).

3. Mizoguchi, T. et al. Distinct Roles of GIGANTEA in Promoting Flowering and Regulating Circadian Rhythms in Arabidopsis. Plant Cell 17, 2255–2270 (2005).

4. Duek, P. D. & Fankhauser, C. HFR1, a putative bHLH transcription factor, mediates both phytochrome A and cryptochrome signalling. The Plant Journal 34, 827–836 (2003).

5. King, R. W., Hisamatsu, T., Goldschmidt, E. E. & Blundell, C. The nature of floral signals in Arabidopsis. I. Photosynthesis and a far-red photoresponse independently regulate flowering by increasing expression of FLOWERING LOCUS T (FT). Journal of Experimental Botany 59, 3811–3820 (2008).

6. Lisson, S. N., Mendham, N. J. & Carberry, P. S. Development of a hemp (Cannabis sativa L.) simulation model 2.The flowering response of two hemp cultivars to photoperiod. Aust. J. Exp. Agric. 40, 413 (2000).

7. Sun, C. et al. Bifunctional regulators of photoperiodic flowering in short day plant rice. Front. Plant Sci. 13, 1044790 (2022).

8. Yano, M. et al. Hd1, a Major Photoperiod Sensitivity Quantitative Trait Locus in Rice, Is Closely Related to the Arabidopsis Flowering Time Gene CONSTANS. Plant Cell 12, 2473– 2483 (2000).

9. Hu, Y. et al. OsPRR37 Alternatively Promotes Heading Date Through Suppressing the Expression of Ghd7 in the Japonica Variety Zhonghua 11 under Natural Long-Day Conditions. Rice 14, 20 (2021).

10. Xue, W. et al. Natural variation in Ghd7 is an important regulator of heading date and yield potential in rice. Nat Genet 40, 761–767 (2008).

11. Du, A. et al. The DTH8-Hd1 Module Mediates Day-Length-Dependent Regulation of Rice Flowering. Molecular Plant 10, 948–961 (2017).

12. Leckie, K. M. et al. Loss of daylength sensitivity by splice site mutation in Cannabis. http://biorxiv.org/lookup/doi/10.1101/2023.03.10.532103 (2023) doi:10.1101/2023.03.10.532103.

13. Izawa, T. Daylength Measurements by Rice Plants in Photoperiodic Short-Day Flowering. in International Review of Cytology vol. 256 191–222 (Elsevier, 2007).

14. Bouché, F., Lobet, G., Tocquin, P. & Périlleux, C. FLOR-ID: an interactive database of flowering-time gene networks in Arabidopsis thaliana. Nucleic Acids Res 44, D1167–D1171 (2016).

15. Pan, G. et al. Genome-wide identification, expression, and sequence analysis of CONSTANS-like gene family in cannabis reveals a potential role in plant flowering time regulation. BMC Plant Biol 21, 142 (2021).

16. Dowling, C. A. et al. A FLOWERING LOCUS T ortholog is associated with photoperiod-insensitive flowering in hemp (Cannabis sativa L.). http://biorxiv.org/lookup/doi/10.1101/2023.04.21.537862 (2023) doi:10.1101/2023.04.21.537862.

17. Ashdown, I. Phytochrome and PSS: ‘Think Beyond Pink’. (2016).

18. Cheng, M.-C., Kathare, P. K., Paik, I. & Huq, E. Phytochrome Signaling Networks. Annu. Rev. Plant Biol. 72, 217–244 (2021).

19. Izawa, T., Oikawa, T., Tokutomi, S., Okuno, K. & Shimamoto, K. Phytochromes confer the photoperiodic control of flowering in rice (a short-day plant). The Plant Journal 22, 391–399 (2000).

20. Franklin, K. A. & Whitelam, G. C. Phytochromes and Shade-avoidance Responses in Plants. Annals of Botany 96, 169–175 (2005).

21. Westmoreland, F. M., Kusuma, P. & Bugbee, B. Cannabis lighting: Decreasing blue photon fraction increases yield but efficacy is more important for cost effective production of cannabinoids. PLoS ONE 16, e0248988 (2021).

22. Magagnini, G., Grassi, G. & Kotiranta, S. The Effect of Light Spectrum on the Morphology and Cannabinoid Content of Cannabis sativa L. Med Cannabis Cannabinoids 1, 19–27 (2018).

